# FITs: Forest of imputation trees for recovering true signals in single-cell open chromatin profiles

**DOI:** 10.1101/2020.08.12.243121

**Authors:** Rachesh Sharma, Neetesh Pandey, Anchal Mongia, Shreya Mishra, Angshul Majumdar, Vibhor Kumar

## Abstract

The advent of single-cell open-chromatin profiling technology has facilitated the analysis of heterogeneity of activity of regulatory regions at single-cell resolution. However, stochasticity and availability of low amount of relevant DNA cause high drop-out rate and noise in single-cell open-chromatin profiles. We introduce here a robust method called as Forest of Imputation Trees (FITs) to recover original signals from highly sparse and noisy single-cell open-chromatin profiles. FITs makes a forest of imputation trees to avoid bias during the restoration of read-count matrices. It resolves the challenging issue of recovering open chromatin signals without blurring out information at genomic sites with cell-type-specific activity. FITs is generalized for wider applicability, especially for highly sparse read-count matrices. The superiority of FITs in recovering signals of minority cells also makes it highly useful for single-cell open-chromatin profile from *in vivo* samples.

First made online as thesis work at https://repository.iiitd.edu.in/xmlui/handle/123456789/807

## Introduction

High-throughput sequencing has enabled a wider application of epigenome profiles for studying biological and clinical samples. Different kinds of epigenome profiles such as histone-modifications (1), chromatin-accessibility and DNA-methylation patterns have been used to study active, poised and repressed regulatory elements in the genome (2). Especially, for characterizing non-coding regulatory regions like enhancers, epigenome profiles have proved to be very useful (3). In the previous decade, epigenome profiling was mostly performed using bulk samples containing millions of cells. Bulk sample epigenome profiles do not help in identifying poorly characterized cell populations and rare cell types in samples of early developmental stages or tumours. Even with *in vitro* experiments, where cells differentiate, there is heterogeneity among single-cells in terms of response to external stimuli. Such heterogeneity is often not captured by using bulk epigenome profile. Studying dynamics of single-cells using epigenome(4) becomes inevitable, especially when single-cell RNA-seq (scRNA-seq) profile cannot explain the difference in response of cells with similar gene expression profile. Such as it was recently shown by Ankur et al.(5) with example of two types of oral cancer cells which had features of epithelial cells and had similarity with each other in terms of overall single-cell transcriptome profile. However, as the drug was applied, one type of oral cancer cells had transition towards epithelial to mesenchymal (EMT) state while other cells remained epithelial. It reveals a non-trivial problem of explaination while relying only on transcriptome profile, and hints towards variation in poising of underlying epigenome. To explain such phenomena, researchers have developed techniques to profile genome-wide epigenome patterns in single-cells. Even though profiling of DNA methylation(6) and histone modification for single-cells is feasible(7), recent large scale single-cell epigenome profiles (8) have been produced using single-cell open-chromatin detection technique(9).

Single-cell open-chromatin profiling can be done using different kinds of protocols like DNase-seq (Dnase I hypersensitive sites sequencing)(10), MNase-seq (Micrococcal-nuclease based hypersensitive sites sequencing)(11) and ATAC-seq (Transposase-Accessible Chromatin using sequencing)(12). Single-cell open-chromatin profile has the potential to reveal both active and poised regulatory sites in a genome. Most importantly, it has recently lead to an understanding of the regulatory action of transcription factors (TFs) when cells are in the state of transition(13). Besides understanding of heterogeneity among cell states, single-cell open chromatin profiles have also proved to be useful for determining chromatin-interaction patterns(14). For analysing single-cell open-chromatin profile, the first step is to do peak-calling after combining reads from multiple cells or using matching bulk samples. Then for each cell, the number of reads lying on the peaks is estimated. While doing so, most often researchers use a large number of peaks, sometimes exceeding more than 100000 in number (8), to capture the signal at cell-type-specific regulatory elements in heterogeneous cell-types. However, due to low sequencing depth and a small amount of genetic material from single-cells, the read-count matrix is often very sparse, which creates a demand for imputation techniques. Using a small number of hyper-active peaks to reduce sparsity may highlight only ubiquitously open sites like insulators and promoters of house-keeping genes which do not have cell-type specificity. Thus with a large number of peaks, single-cell open chromatin profiles have higher chances of including cell-type specific sites but at the cost of a high level of noise and sparsity. The sparsity in the read-count matrix of single-cell open chromatin profile is due to two reasons. The first reason is the high drop-out rate due to which many active genomic sites remain undetected (false zeros). The second reason is the genuine biological phenomenon that there is a large number of silent sites because of their cell-type specific activity. Thus, in comparison to scRNA-seq data, there are higher fractions for both true and false zeros in the read-count matrix of single-cell open chromatin profile. Given such limitations with single-cell open-chromatin profile, the classification and sub-grouping of cells is a difficult task, which is a pre-requisite for many imputation methods.

Due to the reasons mentioned above, most of the imputation methods developed for single-cell RNA-seq (scRNA-seq) profiles, could underperform on single-cell open-chromatin data-sets. Moreover, skipping imputation and doing binarisation for scATAC-seq read-count cannot be fully justified because highly active genomic sites tend to be bound by a large number of TFs. Thus, the binding by a large number of TFs causes wider region with accessible chromatin at highly active sites. Hence even if take unique reads we would have read-count value more than 1 for peaks of size greater than 500 bp. Thus the read-count magnitude is indicative of activity-level of a genomic site. Hence for proper quantification of DNA accessibility using single-cell open-chromatin profiles, there is a need for a second-generation imputation method which can overcome the weakness of other such tools to handle high levels of noise and sparseness. Due to a large number of non-coding sites with cell-type-specific activity, the ideal signal-recovery method must enable detection of such sites like enhancers. Especially with recent droplet-based single-cell ATAC-seq (scATAC-seq) protocol (15), providing profiles of large number of cells with low sequencing depth, the problem of imputation becomes more eminent and challenging.

Even though there has been less attention on imputing scATAC-seq profiles, it is worth noting that many imputation methods have been proposed for scRNA-seq data-sets. MAGIC(16) is the first available method for imputing scRNA-seq profiles. MAGIC predicts missing expression values by sharing information across similar cells, using the approach of heat diffusion. The approach of MAGIC involves creating a Markov transition matrix, constructed by normalizing the similarity scores among single-cells(16). While imputation of expression for a single-cell, the weights for other cells is determined using the transition matrix. MAGIC uses K nearest neighbour (KNN) approach for imputing, however unlike classical KNN based imputation methods(17) it uses a variable value for K. Methods like MAGIC may introduce artefacts into the data and blur out genuine biological variation due to their approach of considering all zero counts as missing values. Another imputation method called scImpute (18) also tries to perform imputation on drop-out genes. For this, scImpute first learns the probability of drop-out for every gene in each cell based on a mixture model for the distribution of read-counts. scImpute predicts the missing values at false zeros by using the information of the same gene in other similar cells which it finds using genes with non-zero expression. Methods like scImpute, which use parametric method to estimate drop-out rate may not be successful for scATAC-seq in estimating true parameters due to inconsistencies in the distribution of tag-counts with very high drop-out rate and noise. High level of sparsity and noise in scATAC-seq profile reduces the chance of finding the correct neighbourhood and sub-clusters of cells, which is an important step for most of the imputation methods like scImpute and MAGIC. Most recently, an approach based on deep-learning called Deep Count Autoencoder (DCA) has been proposed for denoising and imputing single-cell expression profiles. DCA uses an auto-encoder to model and predicts the distribution of the genes using a zero-inflated negative binomial prior. DCA uses auto-encoder to predict the mean, standard deviation and drop-out probability for every gene. For DCA, the mean parameter of the distribution-represents denoised reconstruction. Application of DCA on single-cell open-chromatin profile seems to be a sensible approach. However, single-cell chromatin profiles are much more sparse than single-cell RNA-seq data, hence modelling the distribution of read-counts might not always be successful.

For single-cell open chromatin profiles, we realised the limitations due to improper classification and modelling of the distribution of tag-counts to estimate drop-out rate. Therefore, we developed a method which can overcome these limitations by avoiding sub-optimal solutions, using an ensemble of imputing trees. We call our ensemble-based approach as Forest of imputation trees (FITs). We have benchmarked FITs using scATAC-seq profiles of several cell types using criteria which are useful for analysis for single-cell open-chromatin. Using the scATAC-seq profile of 5 cell types, first, we show that FITs correctly recovers the chromatin accessibility of sites like enhancers with cell-type-specific activity, without performing over-imputation. We also show that FITs is more efficient than other methods in improving dimension reduction and clustering purity for scATAC-seq profiles. Further, we show that unlike other imputation methods, FITs can handle unbalanced scATAC-seq data-sets and helps to avoid detecting false heterogeneity and improves detection of minor cell type. Next, we show that FITs based restoration of read-count matrix also helps in improving prediction in chromatin interaction using scATAC-seq profile.

## Material and Methods

### Pre-processing of Data

We first check the quality of data and remove the peaks which do not have non-zero read-count in any cell. We normalise scATAC-seq read-counts and take log transform of data. Hence the read-count *x*_*ij*_ on a site *g*_*j*_ in cell *i* is represented as

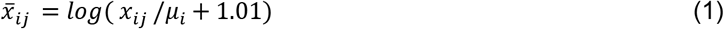

Where *μ*_*i*_ is mean read-count in cell *i*. The log of the normalised matrix of the read-count is provided as input to FITs.

### Clustering using randomized features in a hierarchical manner to improve imputation

Given the noise and sparsity in single-cell open-chromatin profiles finding correct subclasses is not a trivial task. Thus, we use a semi-randomised approach of clustering hand in hand with imputing. Our method has two phases: The first phase consists of making multiple imputed versions of the raw matrix through repeated clustering in hierarchical tree fashion and imputation at each node. Unlike other suggested methods, we do not use all the features at a time to perform clustering as well as we do not perform classification using the raw read-count matrix. We use an iterative approach in every tree such at every parent node we do preliminary imputation followed by classification of cells. The classification is not done only at the bottom nodes (at 3^rd^ layer here).

In the second phase, a final imputed matrix is assembled using the outputs from multiple trees in the first phase.

### Phase-1: The first phase is described below

Given a read-count matrix X to be imputed, where cells are represented by column and peaks by rows, we use the following approach to perform imputation in a tree

Step1: Perform a preliminary Imputation using a base method, over matrix X taking all cells in one class.

Step2: Select n sites(peaks) randomly and perform dimension reduction using t-SNE or singular value decomposition on imputed data. After reducing the dimension, apply k-mean clustering to divide cells among classes. The number of cluster K is randomly chosen within a given range (between 2 - 6).

Step3: After finding classes using K-means clustering, the raw read-count of cells in each class are assembled. After assembling read-count matrix for a class, some peaks appear to have zero read-count in all cells of that class. Hence for imputation on the raw read-count matrix of cells of a class, the peaks with all zero signals are considered as true zeros and dropped.

Step5: The imputed matrix of every class is used further to find sub-classes using the approach mentioned above in step2 and step3. Again, we randomly choose peaks (features) and value for k for the k-mean clustering.

Step6. The non-imputed raw read-count vectors of cells belonging to a subclass are assembled together in a separate matrix. Once again, the peaks which have zero read-count in all cells of a sub-class are dropped and imputation is performed separately for a matrix of each sub-class.

Step 7: The imputed read-count matrix from each sub-class is collected, and a full matrix is built. While doing so, the sites dropped in cells belonging to a subclass are given the value zero. Notice that a version of full matrix is also made using imputation for different classes at first level. Thus from every tree, we collect two versions of the imputed matrix.

The above steps 1-7 are repeated many times to get an ensemble of imputation trees. The output from several trees from phase-1 is further processed in phase-2 using the steps described below

### Phase-2: Following steps are taken in phase 2

For every cell correlations between its unimputed read-count vector and imputed versions from Phase-1 are computed. For every cell average of *m* most correlated imputed versions, is taken as the final imputed vector. For *m* =1, one has to just take top most correlated imputed version.

The reason and logic for some steps in phase-1 and phase-2 are explained below

1. At every node of the tree, initial imputation is done before dimension reduction and clustering, so that chance of getting the correct cluster is high.
2. Further, sub-classification is done so that cells belong to a minor cell-type or cell-state could get chance to come together to have more accurate imputation
3. The imputed version of read-count at level-1 of a tree is also collected so that if a cell belongs to a majority class, we should not force its imputation using smaller groups of cells.
4. In phase-2, we use spearman correlation to choose best k imputed version. We tried several kinds of distance measure to calculate the similarity between unimputed and imputed read-count vectors and found that spearman correlation-based selection of the most suitable imputed version provided the best results.

The step of choosing imputed vectors which have the highest correlation with unimputed read-count is also inspired by the minimisation criteria followed by nearly all imputation methods where the difference between imputed and non-imputed matrices is minimized at observed features, to avoid under and over-imputation. Thus, in other words, we can say that FITs applies the minimisation criteria two times, one during imputation at every node of trees in phase-1 and other at the stage of phase-2.

### The base imputation method of FITS

Even though FITs is designed to be robust to handle error caused during imputation, it is worth describing the underlying base imputation method used by FITs. The base imputation method uses the approach of nuclear norm minimisation with singular value soft-thresholding, as explained below.

Given a read-count matrix Y of a set of cells, where columns represent peaks and rows are for individual cells. The observed read-count matrix Y can be called a sampled version of true ideal matrix *X*. It can be represented as

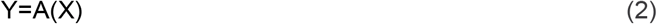

Here *A* is an operator matrix which causes sub-sampling, and has 0’s where the elements of *X* is not observed and 1’s where it is known. The problem of imputation here is to recover complete matrix *X*, given the read-counts in *Y*, and the sub-sampling mask *A*.

Most often approximate rank of the matrix *X* is not known, so getting a solution for equation (2) is not easy. In order to resolve these issues, researchers use an alternative solution. For this purpose, researchers try to solve equation (2) with a constraint that the solution is of low-rank. This mathematical representation for this can be written as,

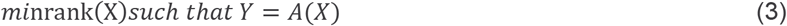

However, this problem itself is NP-Hard. Therefore it’s closest convex surrogate; nuclear norm minimisation is used by many studies(19,20) for matrix completion. The nuclear norm minimisation can be termed as

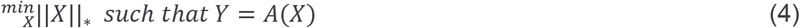

Here ǁ.ǁ_*_ represents the sum of singular values of data matrix X and is called as nuclear-norm. This constraint of minimum rank can be replaced with *l*_1_ norm of the vector of singular values of X as a stringent and convex alternative. Hence as a solution, a modified version of the above equation is proposed(20) as

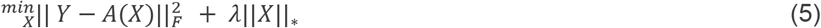

Here λ is the Lagrange multiplier. There is no closed-form solution for the problem in equation (5). Therefore, it is solved in many iterations. To solve such problem, we use majorization-minimization approach at iteration k, given below

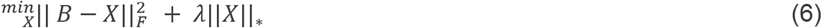

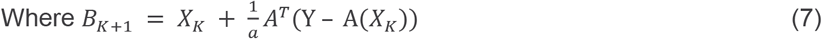

Using the inequality 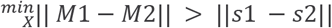 where s1 and s2 are singular values of matrixes M1 and M2, we can express the minimisation problem as

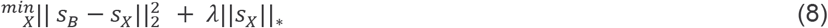

Where *S*_*B*_ and *S*_*X*_ represent singular values of B and X respectively and ǁ*S*_*X*_ǁ is the sum of absolute of singular values of X (21). Thus the minimization problem in equation (7), is often solved by soft thresholding (21) in the following manner

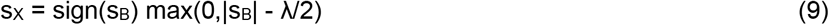

It has been found that the algorithm is robust to the value of λ as long as it is reasonably small (22).

### Estimating co-accessibility among sites and evaluating the prediction of chromatin interaction

Given a read-count matrix of single-cell open-chromatin profile, if we have to find co-accessibility among genomic sites, we can calculate a covariance matrix. However, as the number of elements to be calculated in the covariance matrix is usually larger than the number of data-points in the read-count matrix, estimating the true covariance matrix is not a trivial task. Moreover, the covariance matrix may not always represent direct interaction among genomic sites. Therefore, Graphical Lasso(23) is quite suitable for this kind of problem. The Graphical Lasso method helps in estimating regularized covariance and inverse of the covariance matrix, which can be used to calculate partial correlations between variables (genomic site) (23). The partial correlation represents the measure of the degree of association between two variables when the effect of other variables is removed. Given the noise and small size of data, Graphical Lasso aims to detect a small fraction of true partial correlations among variables. It uses a penalty term which cause shrinkage of partial correlations between many pairs to value zero, if there is not enough strength in the estimate of their association. Graphical Lasso aims to minimize :

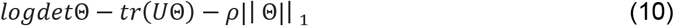

Where Θ is the inverse covariance matrix having the dependence structure of variables and *U* is their covariance matrix and ρ is the penalty term for L1 norm based regularization. Unlike Cicero (by Pliner et al.(24) we did not use the technique of having a penalty term dependent on the distance between genomic sites, as we did not want to miss distal interaction. Moreover, our target here was just to evaluate the improvement in the prediction of co-accessibility by imputation. Here we used the value ρ = 0.01.

Before estimating co-accessibility, we merged peaks which were within 25kb of each other and also added their read-counts. We performed this task on both imputed and non-imputed read-count matrix before calculating their covariance matrix and applying Graphical Lasso. We calculated partial correlation values (co-accessibility scores) between each pair of the genomic region (peak), using the inverse covariance matrix estimated by Graphical Lasso. We downloaded the processed chromatin interaction files for K562 and GM12878 provided by Rao et al. (25) in HiC data format (.hic format). Using files in .hic format, we derived the interaction using Juicer(26) and converted the output to 6 column bed format with scores. Out of all interactions, we chose high confidence interactions with P-value less than 1E-9. We used PGLtool(27) to find overlap between high confidence HiC based chromatin interaction and predicted interacting peak-pairs using co-accessibility.

### Evaluation measures for separability and clustering

After t-SNE (t-distributed Stochastic Neighbor Embedding) (28) based dimension reduction of the imputed and non-imputed read-count matrix, we performed k-means clustering. We used two measures to judge the different properties of clustering and imputation. The first method called adjusted Rand index has cost for false positive and false negatives, where “positive” means that cells of the same type are clustered into one cluster, and “negative” means that two similar cells are assigned different clusters. Let, *T* = [*t*_1_, …, *t*_*P*_] represent the true p classes consisting of *n*_*i*_ number of observations in class *t*_*i*_ and V= [*v*_1_, …, *v*_*K*_] be the clustering result with ‘k’ clusters having *n*_*j*_ number of observations in cluster *v*_*j*_. ARI is calculated as:

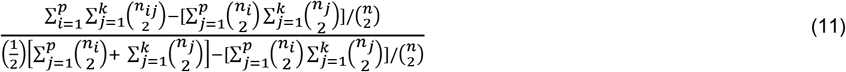

The second measure we used is called as cell type separability (CTS). To calculate CTS, first, we find spearman-correlation between read-counts of each possible cell pair. Then we calculate the median correlation for pairs of cells belonging to the same type to get the intra-cell-type median correlation. Then among two cell-types, we calculate inter-cell-type median correlation by taking only those pairs where one cell belong to one of the 2 types. The difference between intra-cell-type and inter-cell-type median correlations is called as CTS. Thus, CTS value is always calculated between two cell-types.

### Data description and availability

The data-sets used in this manuscript are available in public repositories. The reads for a scATAC-seq profile published by Buenrostro et al. (2015) were downloaded in SRA format (SRA ID: SRP052977). The reads were aligned to hg19 version of the human genome using Bowtie(29). Peaks of ATAC-seq human cell types used here are provided in the same study by Buenrosto et al. at GEO database (GSE65360). The peaks were merged using bedtools. The read-count for every cell was then estimated for peaks in the merged peak list. The scATAC-seq read-count for immune cells was downloaded from GEO database (ID: GSE96772)(30). Single-cell ATAC-seq read-counts for cells from Bonemarrow and liver of adult mouse is available with GEO ID: GSE111586.

For evaluation, we used peak-list of bulk sample ATAC-seq profile of 3 cell types BJ, GM12878, H1-HESC from other GEO database (GEO ID: GSE65360). For defining enhancers, ChIP-seq peaks of histone modification H3K27ac were used. The H3K27ac peak-list for HESC, GM12878 are made available by ENCODE consortium and made available in UCSC genome browser(1). The peak-list for H3K27ac ChIPseq for HL60 is available with GEO IDs: GSM2418804(31). The chromatin interaction files for K562 and GM12878 cell lines downloaded in .hic format have been made available by Rao et al. (2014) (GEO ID: GSE63525) (25).

## Result

Biologically similar cells would have similar activity level at a regulatory site, and this fact can be used to impute the missing values. Hence, if we group similar cells in a sub-cluster, an imputation method has a high probability of providing correct results. However, given the noise, sparsity and imbalance in single-cell open-chromatin data-set, achieving correct sub-cluster is not a trivial task. Hence our method uses randomisation with multiple hierarchical tree-based clustering hand-in-hand with imputations using a base method (Figure 1.). The base imputation method used at every node in the tree uses a known procedure of soft thresholding of singular values for matrix completion (see Methods). The hierarchical tree-based approach used by our method is such that we first perform an initial imputation for all the cells taking them in one group. Using the initial imputed read-count matrix, we classify the cells in into K classes (nodes). For cells belonging to each class, we perform imputation using their raw read count and ignore the previously imputed matrix. However, when we assemble the read count of cells belonging to a particular class, multiple peaks (genomic sites) have zeros read count in all cells belonging to that class. This is exactly as expected, and we utilise it to improve imputation. We consider those peaks with zero read counts in all cells in a class as true-zeros and drop them while doing class-wise imputation. Again, we use the imputed read-count of non-dropped peaks of cells belonging to a class for further classification. Thus, we get subclasses of cells and we again group the raw read count of cells belonging to a subclass. One important point to be noted is that for every level of classification, we randomly choose 50-100% of the non-dropped peaks for classification. We also randomly choose the number of classes k in k-mean clustering at every step. Thus, we perform many such hierarchical tree-based clustering and imputation while randomly deciding k (number of classes) and features for classification. After having final imputed matrices from many such trees, we use the best jth column of multiple imputed version of read-count matrix, for a cell j based on correlation with raw read-count. It is based on our observation that spearman correlation between unimputed read-count vectors of cells of the same type is higher than the correlation between non-similar cells (see supplementary Figure 1). Hence, for a cell, if the imputation is done by clubbing it with wrong neighbours, the imputed vector will have a lower correlation with the unimputed version in comparison to correct imputation. The last step of choosing the best k vectors from multiple imputed version is a crucial filtering step which further helps FITs in avoiding over-imputation (Figure 1). The motivation and logic of different steps of FITs are provided in detail in the methods section.

**Figure 1.**
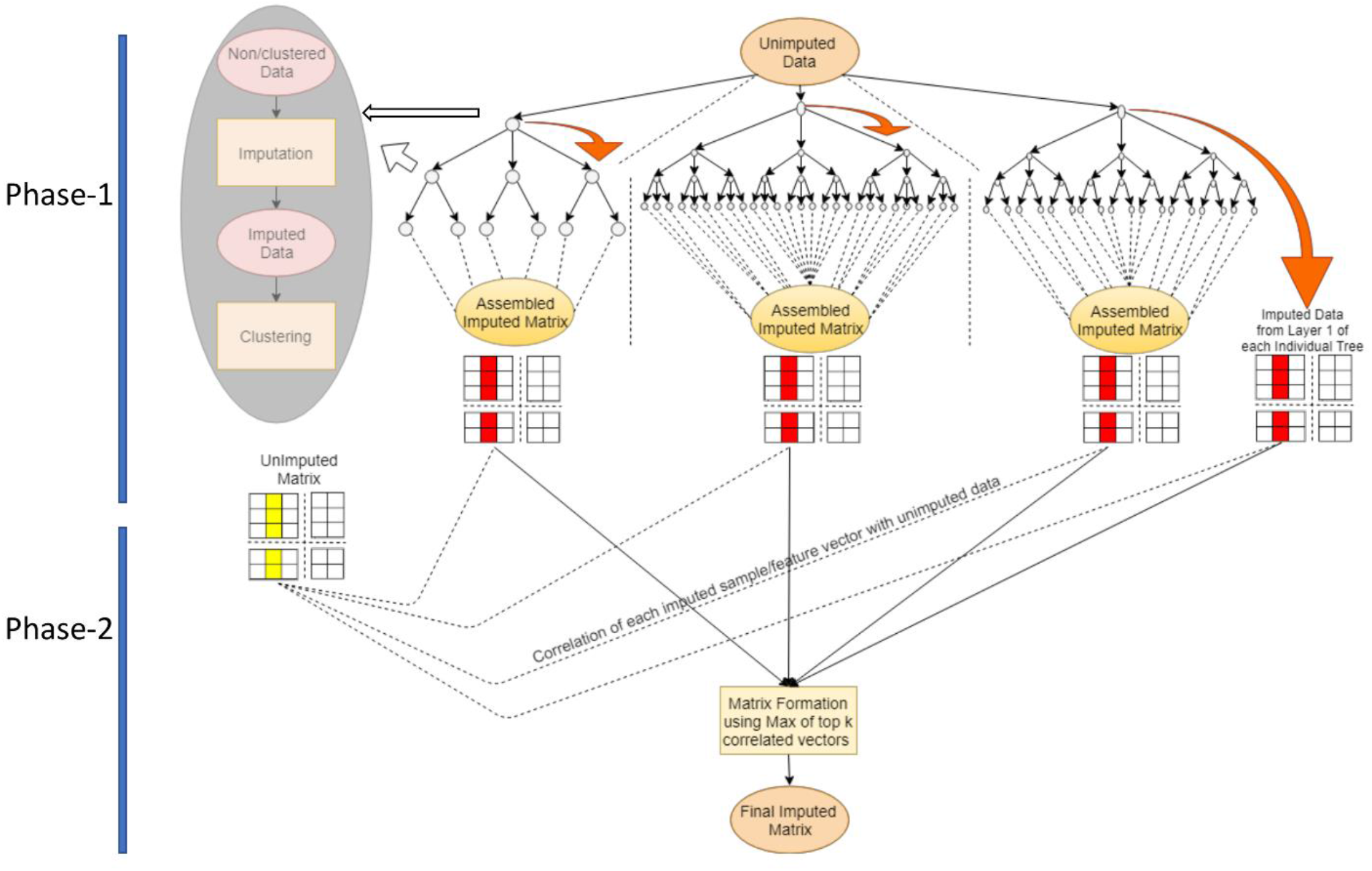
A description of FITs: FITs has two phases. In phase-1 many imputation trees are built to get different imputed versions of the read-count matrix. In an imputation tree, at every node, first, an imputation is performed on the non-imputed read-count matrix using a base method, followed by dimension reduction. Then k-mean clustering is performed to get k clusters of cells. The raw-read count of cells of each cluster is passed on to one daughter node where the same procedure of imputation and further clustering is followed. At every node of an imputation tree, the sites with zeros in all the cells in the raw read-count matrix, are dropped. In phase-2 of FITs, the vectors of multiple versions of imputed matrices are compared to corresponding vectors in the original unimputed matrix using correlation. Finally, for every cell, only those imputed versions are taken which have the highest correlation with its unimputed read-count vector.

### FITs recovers open chromatin signal and avoid over-imputation

We compiled read-count matrix of scATAC-seq profile published by Buenrostro et al. (12) (see methods). Our compiled data-sets had 5 cell types (GM12878, K562, HL60, BJ, HESC) and consisted of 1622 cells and 92447 peaks. We first evaluated if the application of FITs, improves the data quality of single-cell ATAC-seq by correlating it with the relevant bulk ATAC-seq profile. We found that FITs based signal-recovery increased the correlation among bulk and single-cell ATAC-seq profiles (Figure 2A). For B-cell (GM12878) and H1ESC there was almost 4 fold increase in correlation between scATAC-seq and bulk ATAC-seq profiles after application of FITs. Next, we evaluated FITs using promoters of markers genes which are expected to have open-chromatin in a cell-type-specific manner. FITs was able to improve the read-count signal of cell-type-specific promoters without over-imputation in other cell-type. Such as for promoter of CD79a, which has B-cell specific expression(32), FITs caused amplification of its read-count signal only in GM12878 (Figure 2B). Similarly, for the promoter of the SOX2 gene, the imputation by FITs caused an increase in read-count value only for HESC (Figure 2C).

**Figure 2:**
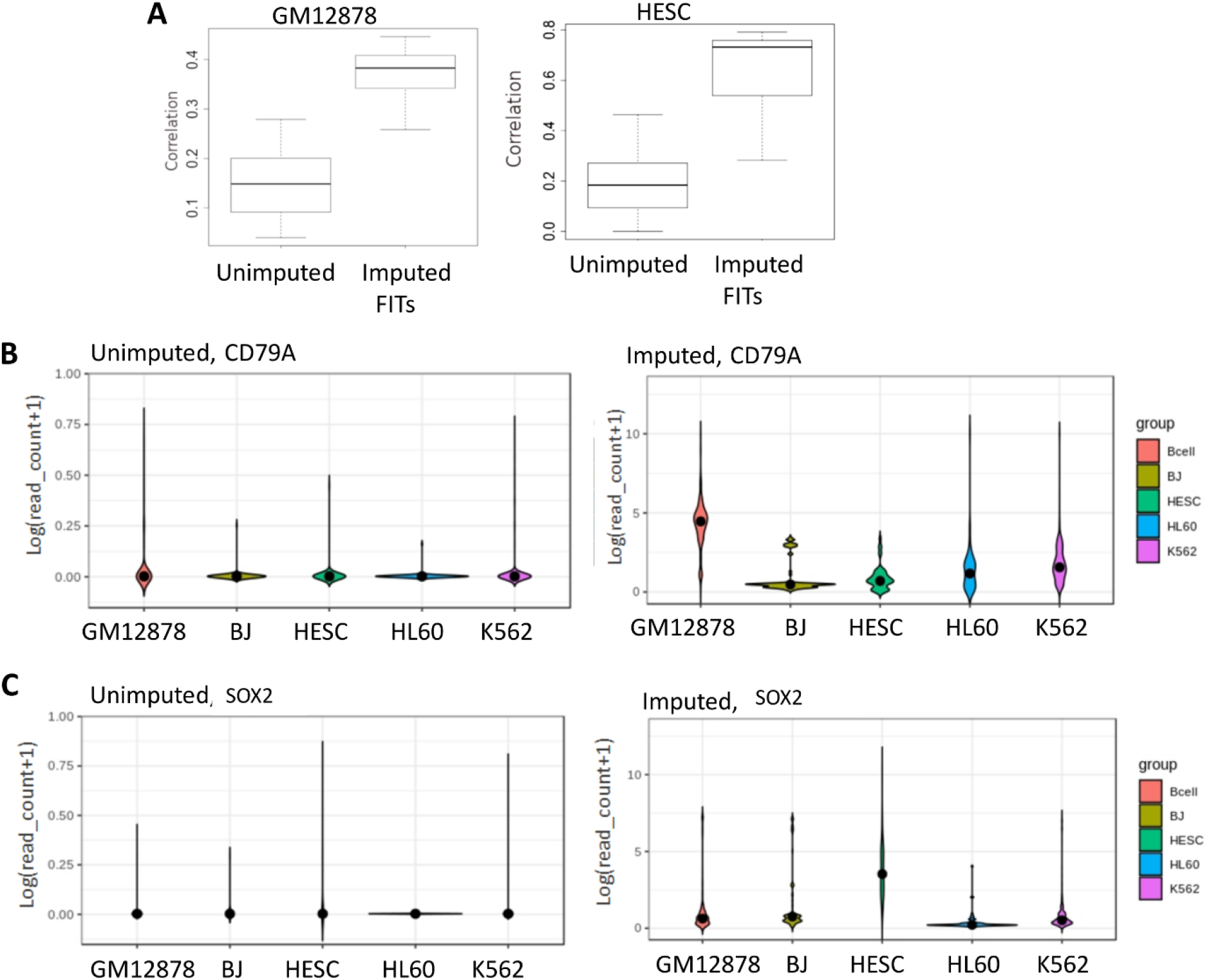
FITs improves signal in single-cell ATAC-seq profile. (A) Boxplot of correlations of imputed and non-imputed scATACseq read-count with bulk ATAC-seq profile in relevant cell-types from other studies. Results for two cell types, GM12878 and HESC are shown here. (B) Violin plot of imputed and non-imputed read-count on the promoter of CD79a which is known to have expression more specifically in B-cells (GM12878 here). (C) Violin plot of read-counts on the promoter of SOX2 gene. Among 5 cell types in our data-set, SOX2 gene is supposed to have expression only in HESC. FITs based imputation shows high read-count of SOX2 only in HESC cells.

We further evaluated the performance of FITs in comparison to KNNImpute, and 3 other methods (MAGIC, scImpute and DCA) developed for single-cell RNA-seq read-count matrices. For every genomic site in the used scATAC-seq data-set, we first found in which of the 5 cell types it overlapped with a genuine peak of bulk ATAC-seq profile. We estimated the coverage for peaks of bulk ATACs-seq in respective cell-types and calculated ROC-AUC for every cell (Figure 3A). FITs based signal-recovery resulted in consistently higher median AUC than other imputation methods for coverage for true peaks from bulk samples.

**Figure 3.**
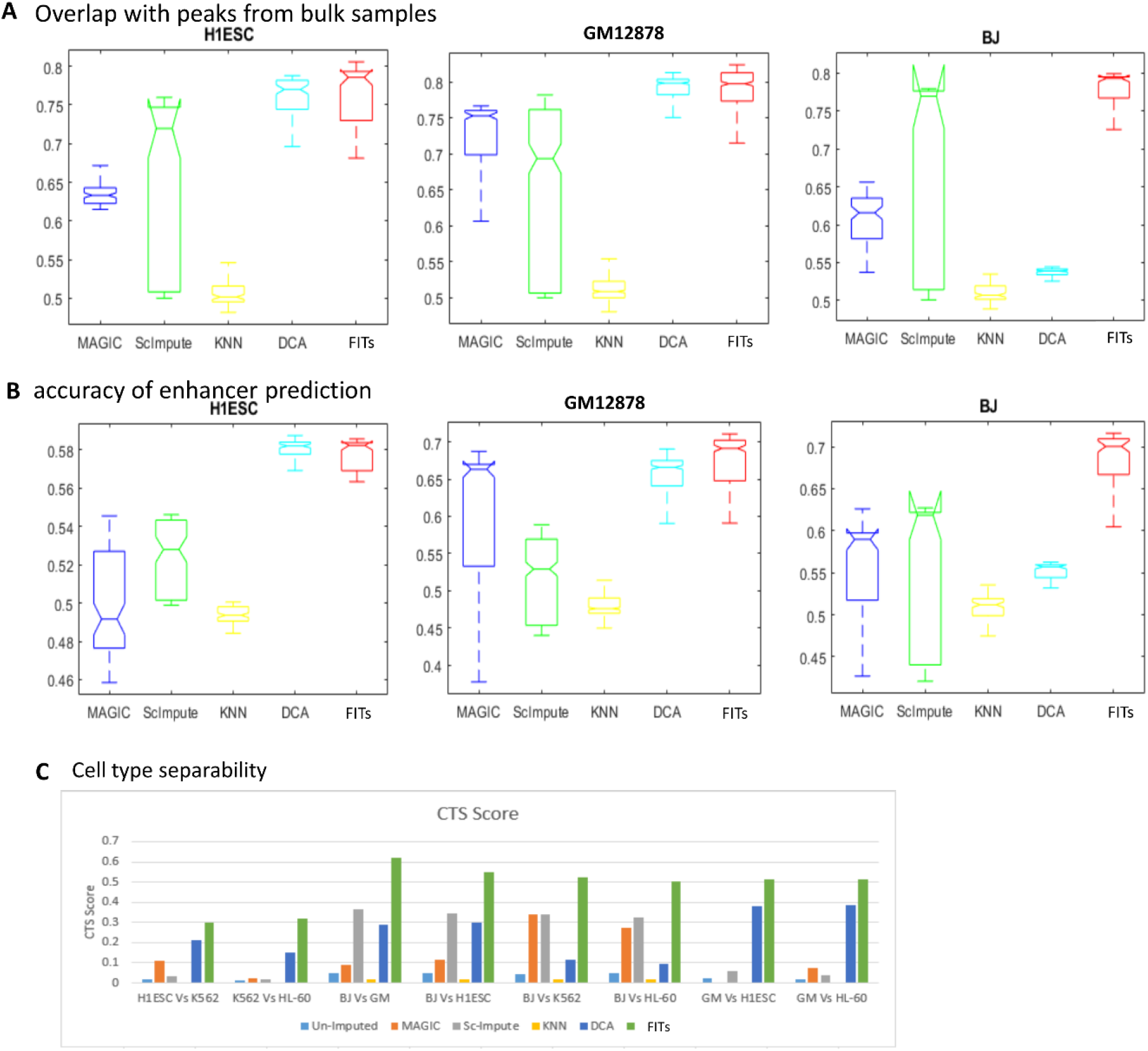
FITs based imputation improves coverage of true peaks and enhancers. For evaluation single-cell ATAC-seq data-set published by Buenrostro et al. (2015) was used here. (A) For every cell coverage for relevant bulk ATAC-seq peak according to intensity of read-count in the imputed read-count matrix was calculated using the approach of ROC (receiver operating characteristic curve). Here positive value means that a site has a peak and negative represent no-peak in relevant bulk ATAC-seq profile of the relevant cell. The area under the ROC (AUC) for each single-cell was calculated and box plots of AUC of cells for different cell types are shown here. (B) Boxplot of AUC for coverage of enhancers using normalised read-count of scATAC-seq profile. The true set of enhancers for a cell-type was compiled using H3K27ac ChIP-seq profile from the bulk sample. (C) Cell type separability (CTS) among cells of different cell-types calculated using imputed read-count matrices.

### FITs improves detection of cell-type-specific sites

One of the main purposes of open chromatin profiling is to study the activity of cell-type-specific regulatory elements like enhancers(33). Few scientific groups have used the technique of highlighting cell-type-specific activity using open chromatin signal to predict enhancers (10). We used a similar technique and divided scATAC-seq read-counts on a peak by its average read-count across all the cells. For validation, we used non-promoter peaks of H3K27ac ChIP-seq profiles of bulk samples of cell-lines, as enhancers(1,31). Further evaluation and comparison with the other four methods revealed that imputation with FITs consistently provided higher coverage for enhancers compared to other methods (Figure 3B). DCA had a comparable performance for H1ESC cells; however, for BJ cells, DCA seems to have failed in recovering a genuine signal (Figure 3B). The performance of MAGIC varied for different cell types, whereas median AUC for scImpute for detection of enhancers remained low in the range of 0.52-0.62 (Figure 3B).

Overall FITs restores signal at cell-type-specific sites in scATAC-seq profiles without over-imputing, which can help researchers to detect enhancers for downstream analysis. The capacity of improving signal at cell-type-specific sites can help in improving cell-type-separability (CTS). Here, CTS is defined as the difference between the median intra-cell-type correlation and inter-cell-type correlation (see Methods). We calculated CTS score among different pairs of cell types (BJ Vs GM12878; BJ vs HESC and GM12878 Vs HESC) and found that FITs based imputation provided the best CTS among all 4 methods used for comparison (Figure 3C). Among other methods, DCA appeared to be second best, but the CTS values for DCA were substantially lower than FITs.

### FITs improves dimension reduction and clustering of single-cell ATAC-seq profiles

One of the major tasks in the analysis of single-cell open-chromatin profile is to reduce the dimension of read-count matrix for visualisation and classification. Due to the high level of noise and sparsity in scATAseq read-count matrix, researchers often resort to calculating accessibility score for transcription factor motifs for dimension reduction based visualisation and classification(8). However, dimension reduction and classification of read-counts directly could reveal new classes and states of cells which could be blurred out by using motif accessibility scores. We performed t-SNE (28) based dimension reduction and visualisation of scATAC-seq read-count matrix for 5 cell lines published by Buenrostro et al. (2015)(12). As expected, the application of t-SNE on raw (unimputed) read-count did not provide satisfactory results as cells of different types were co-localised together in lower-dimensional space (Figure 4A). Similarly, applying t-SNE on read-count matrix imputed by other tools methods also provided results which had a mixing of coordinates for cells of different types. However, with read-count matrix imputed by FITs, the coordinates provided by t-SNE had clear separability among different cell types. It is quite evident from t-SNE plots of imputed matrixes that MAGIC and scImpute introduce artefactual grouping of cells (Figure 4A). MAGIC and scImpute outputs had artefacts possibly due to complete reliability on one-time grouping and sub-classification of the raw read-count matrix for imputation. In the output based on KNNimpute, GM12878 and K562 cells appeared to have overlapping locations in t-SNE based visualisation. On the other hand, DCA seems to have mixed the profile of H1ESC and GM12878 during imputation (Figure 4A). Further, we compared the accuracy of clustering using the imputed scATAC-seq profiles. For this purpose, we used the adjusted Rand index (ARI) after applying K-means clustering on t-SNE based coordinates for imputed read-count matrixes. FITs had highest ARI score among the tested methods. The ARI scores for output of other methods were 2-3 times lower than FITs.

**Figure 4.**
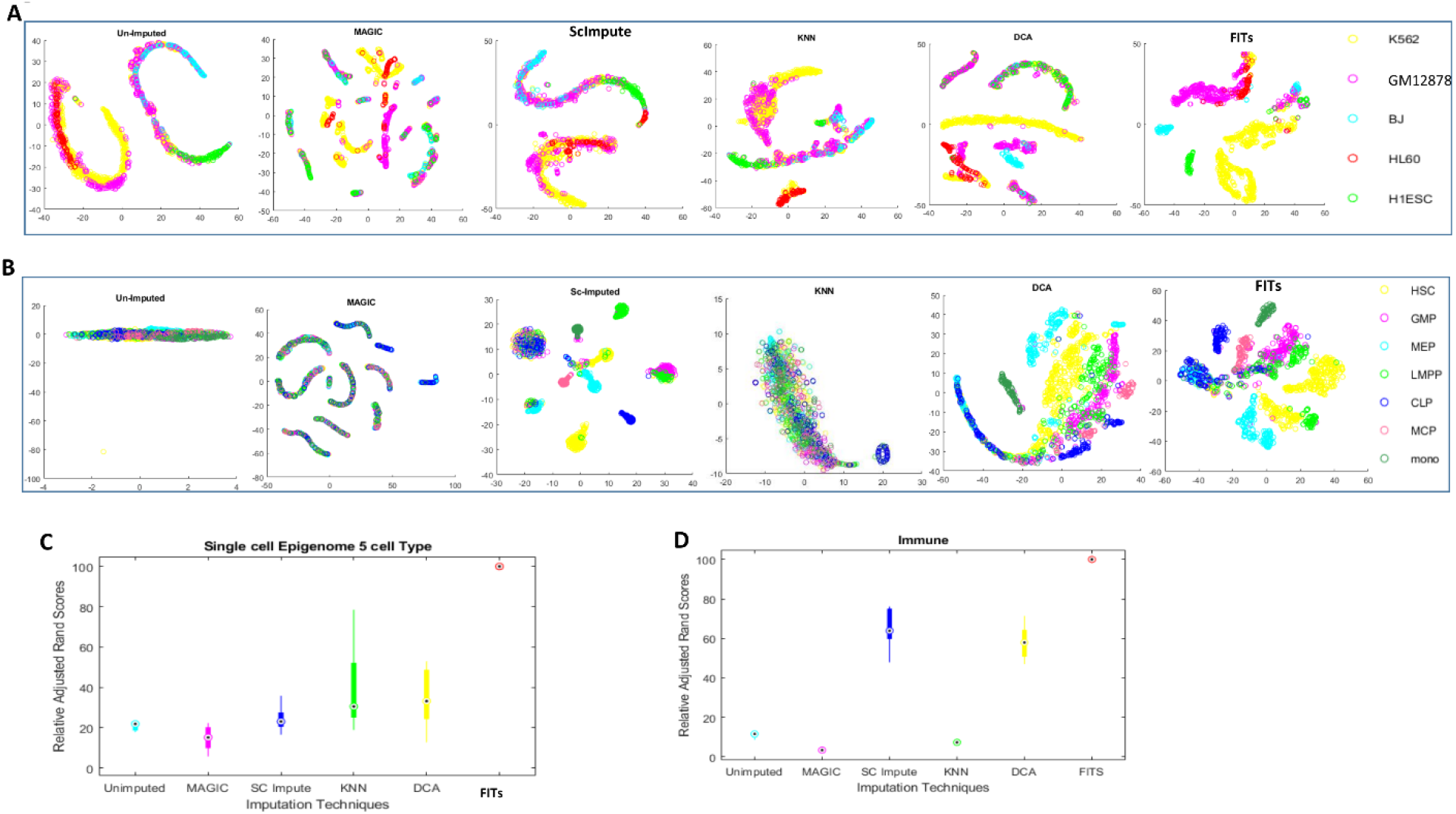
Performance of different imputation methods based on t-SNE based dimension reduction. (A) Scatter plot of result of t-SNE for outputs from different imputation methods for Single-cell ATAC-seq profile from Buenrostro et al. (2015) (B) Visualisation of t-SNE results for unimputed and imputed read-count matrices for hematopoietic cell data-set. (C) Boxplot of Relative adjusted rand index (ARI) to enumerate clustering quality when k-means clustering is applied on t-SNE based reduced dimension. The results clustering of ATAC-seq profile from Buenrostro et al. (2015) is shown here. To make boxplot, ARI was calculated at different values of k in the range of 3 to 10. (D) Boxplot of Adjusted Rand Index (ARI) relative to FITs for evaluation of classification of hematopoietic cells. The relative ARI values were calculated at different values of k (between 5-12) for k-mean clustering.

We simulated higher drop-out rate using the same scATAC-seq data-set of 5 cell lines (by Buenrostro et al.(2015) and randomly dropped 10-30% of read-counts. At all simulated additional drop-out level FITs output provided clear separability among different cell-types in t-SNE based visualisation (supplementary Figure 2 A-C). With a higher drop-out rate in 5 cell-line data-sets, the performance gap between FITs and other methods further increased in terms of clustering purity achieved after imputation. As can be seen in Supplementary Figure 2 D the ARI score for FITS output is 3-4 times greater than all other methods compared for simulated drop-out. Further, we used a data-set which had a severe problem of drop-out as well as overlap among the activity of regulatory sites among different cells. We used scATAC-seq data-set, generated using phenotypically defined human hematopoietic cells, including early hematopoietic progenitors and cells of myeloid and lymphoid lineage(30). This data-set of immune cells had cell-type labels for every cell. After applying different imputation methods followed by t-SNE based visualisation, we achieved similar results such that FITs had more separability among non-similar cell-types (Figure 4B).

Whereas, unimputed version and outputs of MAGIC and KNNimpute had completely mixed co-localisation for different cell types. In terms of ARI even scImpute and DCA were not comparable to FITs (Figure 4D). On careful observation, we found that there are 2 cell types (HSC and LMPP) which have two groups of cells in t-SNE plot for FITs (Figure 4B). Our investigation revealed that those groups came from different batches which might have different culture-microenvironment or experimental setup. Thus, the process of restoration of open-chromatin signal and reduction in noise by FITs improved clustering and separability of different cell-states and batches.

### FITs can handle unbalanced read-count matrices and restore signal of minor cell population

The approach of hierarchical steps of imputation and clustering hand-in-hand with randomisation in FITs has the potential to highlight minor population cluster even in the presence of dominating signal from major cell-types. In order to evaluate the performance of imputation methods on imbalanced data-set of scATAC-seq profile, we first created a data-set consisting of K562 cells with 95% frequency and H1ESC with 5% occurrence rate. After imputation of the simulated data-set, we found that FITs was able to recover the signal of most of the cells of minor cell-type (H1ESC) such that they could localise as a separate group in t-SNE based visualisation (Supplementary Figure 3). Whereas in the case of other imputation methods, H1ESC co-localised with major cell-type (K562) in t-SNE plot of imputed data-set (Supplementary Figure 3). The output of FITs also had highest ARI score for clustering purity of imputed imbalanced data-set.

After evaluating imputation methods for simulated imbalanced data-set, we used in-vivo data-sets for further testing. We performed imputation on scATAC-seq profiles of cells from adult mouse bone marrow and liver published by Cusanovich et al. (8). For scATAC-seq data-set of adult mice, Cusanovish et al. have performed annotation and assigned cell-type for most of the cells. The compiled scATAC-seq data-set for bone marrow had 4033 cell and liver data had 6167 cells. After imputation, we performed t-SNE based dimension reduction and visualization using 8 most frequent cell-types and cells with a label of “unknown”, for both data-sets. Even among retained cell-types, there was an imbalance in numbers. In scATAC-seq data-set of bone marrow, there were 4 cell types (Macrophages, B-cells, Dendritic cells and T-cells) with a frequency less than 2% and two cell types represented more than 70% cells (Hematopoietic progenitor + erythroblast). In liver scATAC-seq data, there was an even higher imbalance in numbers for cells of different types. Among the retained cells in liver scATAC-seq data, 91.4% were hepatocytes, while 6 cell-types had frequency less than 1.6%.

Visualisation of t-SNE results for bone marrow data-sets revealed other tested imputation methods were inefficient in recovering signal of minor cell-types (Supplementary Figure 4). For MAGIC, scImpute and KNNImpute, minor cell type locations got mixed with major cell-types in t-SNE plots. For bone marrow scATAC-seq profile, the imputation by scImpute caused the formation of many clusters within major cell types such as erythroblast, hematopoietic progenitors and monocytes. On the other hand, minor cell types in bone marrow data such as B cells, dendritic cells could not be isolated as separate groups in t-SNE results for matrix imputed by scImpute. MAGIC also had similar results like scImpute in terms of separability for minor cell types. However, for bone marrow data, FITs was efficient in recovering signal even for minor cell types. The t-SNE plots for read-count matrix imputed by FITs showed separability of minor populations cells, such as macrophages formed a group which was clearly visible as a separate group from other cell-types. We used CTS score to evaluate the separability of minor cell types after imputation. For minor cell types scImpute, DCA had noticeable CTS scores, however, they were approximately 1.5 times lower than corresponding CTS values for FITs based output (Supplementary Figure 4B).

Results for imputation of liver scATAC-seq profiles also revealed that when the read-count matrix was highly imbalanced, other methods (scImpute, MAGIC, KNNImpute and DCA) failed completely and could not help in segregating minor cells in t-SNE based visualisation (Figure 6A). On the other hand, FITs based imputation caused the formation of separate groups for minor cell-types. There was a substantial difference between FITs and other 4 tested methods in terms of CTS (Figure 6B) and ARI score (Supplementary Figure 5). Thus, the examples of bonemarrow and liver data-sets, highlight the importance of sub-clustering using tree-based approach in recovering signals of minor cells.

**Figure 5.**
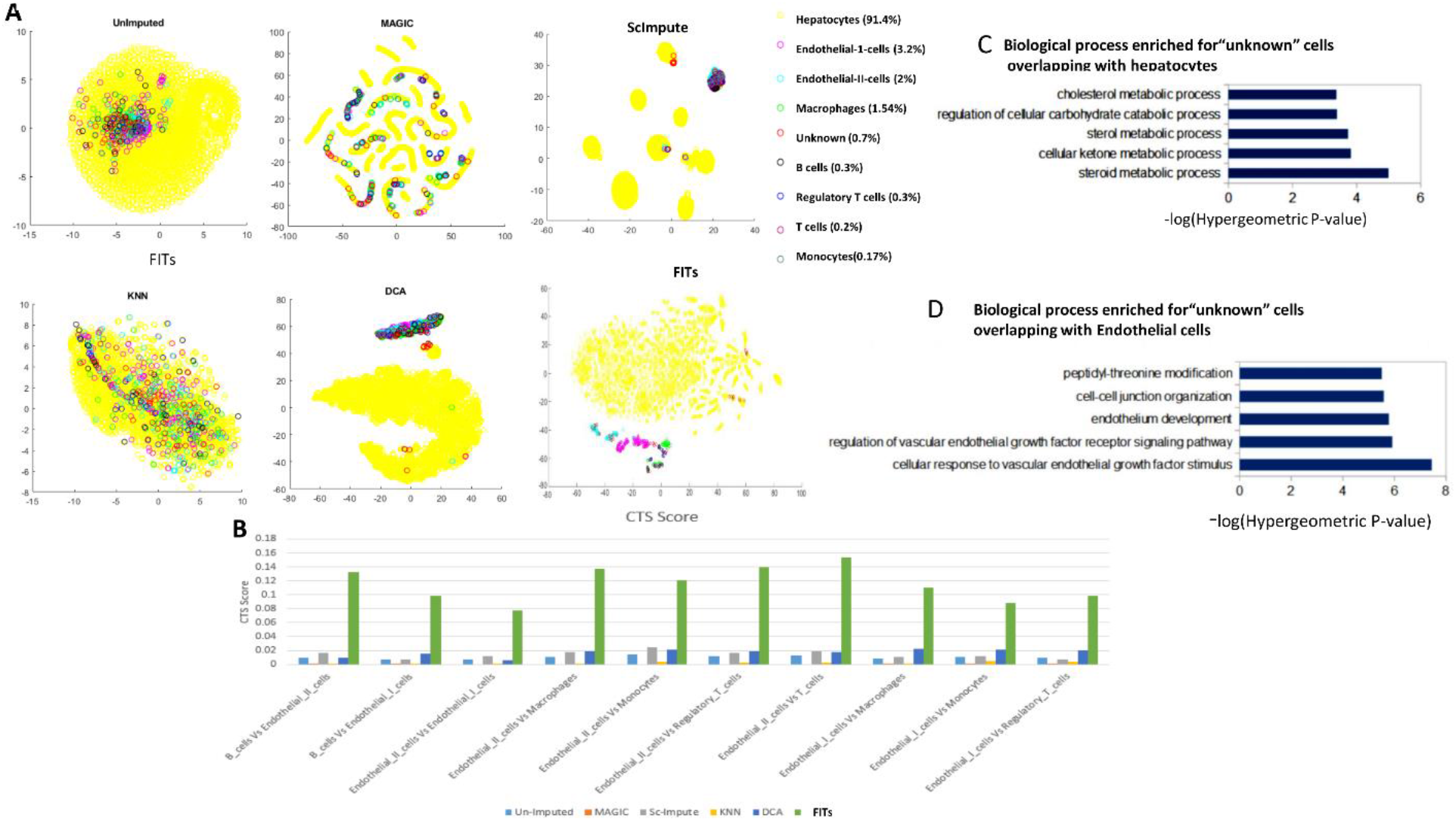
FITs recovers signal of minor cell-types in imbalanced single-cell ATAC-seq profile from *in vivo* sample. The data-set used here is the scATAC-seq read-count matrix for cells in adult mouse liver. (A) scatter plot of t-SNE results for unimputed (raw) and read-counts matrix imputed by 5 methods (B) Cell type separability(CTS) among different cell-types in read-count matrix imputed by 5 methods (C) Top enriched gene-ontology terms (Biological Process) for predicted enhancers of “positive” cells co-localising with hepatocytes in t-SNE based plot for FITs output. For unbiased analysis, enhancers were predicted using unimputed read-counts of unknown cells co-localising with hepatocytes. (D) Top biological process terms enriched for predicted enhancers in unknown cells co-localising with endothelial-1 cells in results of FITs based t-SNE plot. Again, only unimputed read-count were used to predict enhancers.

**Figure 6.**
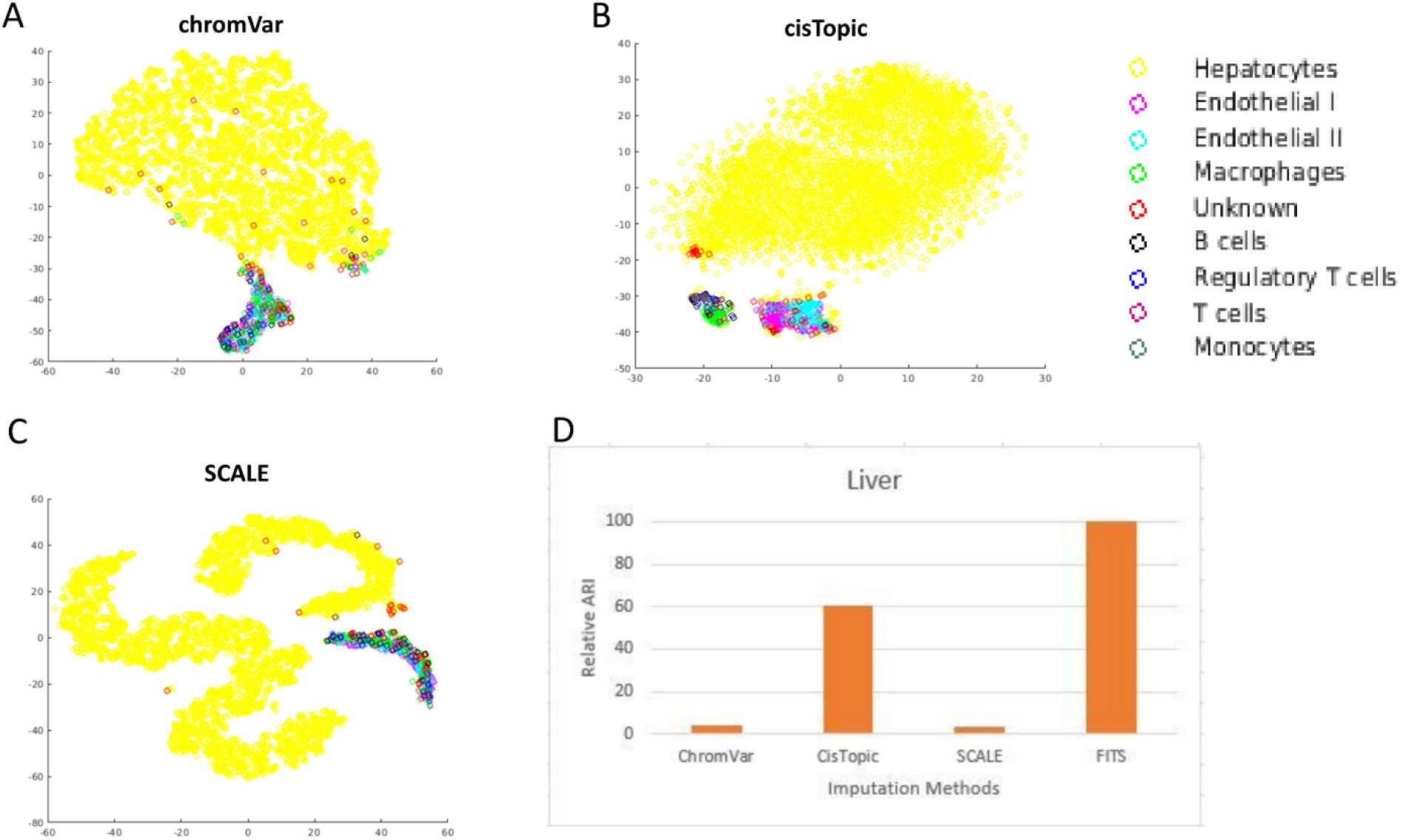
Evaluation of tools designed for dimension-reduction based visualisation of scATAC-seq profile. (A) Visualisation scATAC-seq profile of cells from mouse liver by chromVar (B) by cisTopic (C) by SCALE. (D) Clustering purity in terms of Relative adjusted Rand Index (ARI) with respect to FITs based results. ARI was calculated after dropping the Hepatocytes to assess the recovery of signal of minor cell-types and classification purity for imbalanced data-sets. Here the value of k is 8 for kmeans clustering.

For FITs based imputed version of the liver data-set, few cells with the label as “unknown”, co-localised with hepatocytes in t-SNE based scatter plot (Figure 6A). We normalised the raw read-counts to highlight enhancers in “unknown” cells overlapping hepatocytes. We chose the top 10000 potential enhancers based on the average of normalised read-count for “unknown” cells co-localising with hepatocytes and performed GREAT based gene-ontology analysis(34). The top biological Process terms using GREAT based analysis were related to functions of liver cells such as cholesterol and ketone metabolic processes. Thus it became quite evident that “unknown” cells overlapping with hepatocytes were also typical liver cells. (Figure 6A, C). Using the same procedure on “unknown” overlapping with endothelial-1 cells revealed top enriched gene ontology term related to endothelium development (Figure 6D). Thus, FITs based imputation also enabled annotation of cells labelled as “unknown”.

Most of the previous tools meant for dimension reduction of scATAC-seq profile do not use imputation however for completeness of our study, we compared the performance of 3 such tools (ChromVar(35), cisTopic(36) and SCALE(37) with FITs. With liver scATAC-seq profile, none of the 3 tested tools (chromVar, cisTopic and SCALE) could display minority cell-types separately like FITs based results (Figure 6). Further, we found that computation-time needed by FITs is similar to cisTopic and SCALE (Supplementary Table-1).

### FITs improves detection of chromatin interaction from single-cell open chromatin profile

After evaluating FITs for improvement in calculating similarity among cells, we investigated whether imputation can help in estimating co-accessibility among sites. Recently, Pliner et al. proposed that regions with high co-accessibility in single-cell open chromatin profile are highly likely to be interacting(24). Chromatin interaction maps help in multiple processes such as identification of the target genes of noncoding genomic loci highlighted by GWAS (38) (Genome-wide association study) and understanding of gene regulation. Pliner et al. applied Graphical Lasso(23) based approach to predict interaction among genomic sites(24). Even though the graphical Lasso method(23) is used to reduce the effect of noise and to calculate direct interactions, it’s performance could be improved by providing less sparse data. We applied graphical Lasso based approach to evaluate imputation based improvement in the prediction of chromatin interaction using scATAC-seq. We used HiC based chromatin interaction profile for K562 and GM12878 published by Rao et al. for evaluation(25). Using hic files we extracted high-confidence chromatin interactions at 25Kbp resolution for both K562 and GM12878 cell lines. For both K562 and GM12878 cells, we merged the peaks lying within 25 Kbp in scATAC-seq read-counts matrix and applied Graphical Lasso to detect intrachromosomal chromatin interactions. For both cell types K562 and GM12878, FITs based imputation indeed improved overlap among predicted and true high-confidence chromatin interaction by 10-30% for different chromosomes (Figure 7). One important issue to be noticed is that unlike Cicero, we did not focus on predicting interaction only within a certain distance range. Rather we also predicted intra-chromosome interaction between sites lying far apart. Thus, FITs prove to be useful for analysis of single-cell open-chromatin profile in multiple different ways, including chromatin interaction prediction.

**Figure 7.**
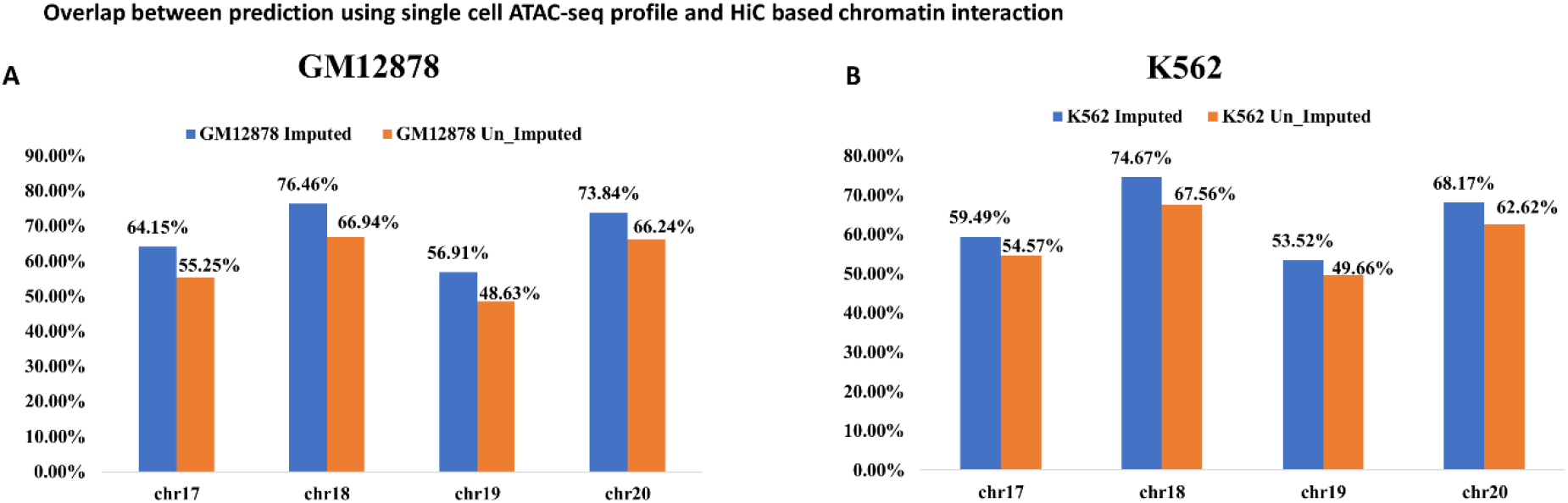
FITs based imputation improves prediction of chromatin-interaction using scATAC-seq profile. The fraction of co-accessibility based predicted interaction overlapping with chromatin-interaction from HiC, is shown here. (A) for GM12878 cells (B) for K562 cells.

## Discussion

The patterns of signal and sparsity in single-cell open chromatin are different in comparison to RNA-seq and DNA methylation profiles. Therefore, imputation of scATAC-seq profiles need attention and it cannot be treated just like scRNA-seq data-set. Single-cell open chromatin profiles are often used to get insights about the effect of pathways and master regulators through non-coding regulatory sites with cell-type-specific activity. Hence cell-type specificity is a major concern while analysing single-cell open-chromatin profiles. Moreover, there is a need to study the impact of imputation in different kinds of downstream analysis steps for single-cell open-chromatin profile. Therefore, first, we evaluated whether imputation of scATAC-seq can help in improving down-stream analysis desired for single-cell open-chromatin profile. Second, we developed a method for imputing which can handle a high level of noise, sparseness and cell-composition bias in single-cell open chromatin profiles. The strength of FITs lies in 3 features: randomised sub-clustering and imputation in multiple trees to avoid sub-optimal solution, deciding drop-out after clustering and choosing imputed vectors based on correlation with the unimputed version to avoid wrong-imputation.

We have shown here that methods relying on parameters for single clustering step increase chances of artefacts due to errors in classification. Relying completely on few non-randomized classification steps also creates the risk of getting trapped in local minima and failure to detect true heterogeneity. We have shown here that FITs performs imputation in such a way that for scATAC-seq profile, there is less chance of detecting false heterogeneity in comparison to other imputation methods. Especially, when we have an unbalanced data-set, classification often fails to identify the minority population as a separate class, which creates artefacts during imputation by methods like scImpute and MAGIC. There have been multiple studies related to detecting rare cell-states using scRNA-seq profiles; however, with scATAC-seq such analysis is rarely done due to overwhelming noise and imbalance in data-sets. Our analysis using FITs, hints that detecting rare cell-states using scATAC-seq read-counts is feasible, and it can provide a new direction in the analysis of clinical *in vivo* samples. In addition to the recovery of minor-cell signals, we also showed the applicability of FITs for 3 analysis steps peculiar to scATAC-seq profiles which are; enhancer-detection, and chromatin interaction prediction. Thus FITs can partner with many existing tools for improved inference and novel applications of scATAC-seq.

An advantage with FITs is that it can handle huge read-count matrices because of horizontal scalability. To run Phase-1 of FITs, one can break down the huge read-count matrix into many smaller matrices with randomly chosen cells. Two smaller matrices can have the same cell, but the union of all small matrices should represent all the cells in the original data-set. The Phase-1 of FITs can be run for multiple small matrices on different computers, before final matrix compilation by in Phase-2. Other imputation methods are rarely designed to handle very big read-count matrices. Hence FITs also resolves the problem of imputing large read-count matrices.

Multiomics studies using single-cell profile provide a global perspective of development and disease; however, very few groups have made such attempts(39). Thus, the major advantage of FITs is that it would encourage more researchers to explore single-cell open-chromatin profiles for multiomics studies, due to reliability it adds during analysis. Other types of single-cell epigenome profiles such as histone modifications, MNAse-seq and DNAse-seq have also been used in few studies. The generality of FITs makes it suitable for other kinds of single-cell epigenome data-sets also, however in future FITs could be further adapted for other kinds of single-cell epigenome profiles also.

**The Python and Matlab version of FITs and https://reggenlab.github.io/FITs/ and imputed matrices used here for figures can be downloaded from** http://reggen.iiitd.edu.in:1207/FITS/imputed_finaldata/

## Authors contribution

VK and AM designed the study and reviewed the results and the manuscript. AM, Mongia, RS implemented the code in Matlab. SM implemented the code in python. VK arranged read-count for tested data-sets and wrote the manuscript. RS executed the programs in different data-sets and created a few figures. NP made a few figures and analysed chromatin interaction profiles.

## Material and Correspondence

Emails : Vibhor Kumar –, Anghshul Majumdar –

